# Modal properties of fruit-rachilla system of the macaw palm

**DOI:** 10.1101/2020.07.27.222687

**Authors:** Flora Maria de Melo Villar, Francisco de Assis de Carvalho Pinto, Fabio Lucio Santos, Daniel Marçal de Queiroz, Mariana Ribeiro Pereira, Domingos Sárvio Magalhães Valente

## Abstract

The macaw palm has been domesticated due to its potential use in the production of biodiesel. One of the current challenges in this area is to determine the proper information and develop the technologies required for using macaw palm, allowing it to contribute to the sustainable production of feedstock for biodiesel industry. The principle of mechanical vibration can be employed for the detachment of fruits from a tree bunch, and it is therefore necessary to study the dynamic behavior of the macaw palm fruit-rachilla system during the vibration. Hence, the modal properties of the system were determined. A study on the dynamic behaviors was carried out using a deterministic finite element model, and the natural frequencies were obtained through a frequency-scanning test to evaluate the model. The mean relative error (MRE) between the measured and simulated natural frequencies was also used to evaluate the model. The natural frequencies, determined experimentally, varied between 26.21 and 33.45 Hz on average, whereas the simulated frequencies varied from 24.81 to 39.27 Hz. The overall MRE was 9.08%. Once the model was validated, a sensibility test showed that the density of fruit and the elasticity modulus are the parameters that most influence the natural frequencies of the fruit-rachilla system.

## INTRODUCTION

One of the main challenges in the Brazilian agro-energy sector is related to the generation of information and technology to develop a sustainable source of feedstock for the production of biofuels (Lorenzi et al., 2011). The macaw palm is a promising crop as a feedstock for biofuels owing to its high oil yield and quality; however, it is still employed using an extractive system. The use of new technologies for fruit harvesting and processing is essential to allow the macaw palm to become commercially profitable to farmers, and to facilitate intensive exploitation. The oil quality can be affected if the fruit is not harvested and processed within the proper time and through the correct method. Evaristo et al. (2016) observed that the acidity level and oxidative stability of macaw oil are affected by the age of the fruit and its storage period.

Mechanical vibrations can be used as an option for the process of removing the fruit from the rachilla, either in a biofuel plant or during harvesting. Like other fruits, such as coffee (Coelho et al., 2015), the modal properties, natural frequencies, and methods of vibration of the fruit-rachilla system are the basic data required to develop equipment employing this phenomenon.

Because the macaw palm domestication process started recently, a natural and significant variety is present among the different accessions, which generate fruits of different sizes and maturation. To help in the development of machines that can harvest the fruit, it is a general practice to generate a mathematical model of the system, in order to obtain information about its dynamic behavior when subjected to mechanical vibrations. Once we have a proper model, we can simulate the dynamic behavior of the fruit-rachilla system to estimate the modal properties.

The finite element method can be used to solve different problems by dividing the model into the specific elements of the body being studied, and studied, and generating a mesh and a group of equations that describe the behavior of the variables involved (Silva et al., 2014). In the case of coffee, for example, a mathematical simulation using the finite element method was applied to obtain information regarding the modal parameters of the fruit-peduncle (Coelho et al., 2015).

The objective of this work was to develop a model to describe the dynamic behavior of the macaw palm fruit-rachilla system, and to simulate this system to identify which parameter has the greatest influence on its natural frequencies.

## MATERIAL AND METHODS

The fruit bunches used to determine the modal properties of the fruit-rachilla system were collected at UFV’s Active Germplasm Bank (AGB), Araponga city, Minas Gerais state, Brazil. Tests were conducted using the bunches with green fruits. The harvesting of the fruit bunches was performed on Sept. 30, 2015, and the tests were performed on the same date. Four palm trees of different accessions were used and identified as BD 27 (from the Abaeté region, Minas Gerais state), BD 40 (from the Pitangui–Martinho Campos region, Minas Gerais state), BGP 29 (from the Prudente de Morais–Matozinhos region, Minas Gerais state), and BGP 35 (from the Mirandópolis region, São Paulo state).

The mechanical, physical, and geometric properties were used as the input parameters during the computational modeling and simulation of the fruit-rachilla system to determine the natural frequencies and vibration modes (Table 1). The mechanical properties of the fruits, the elasticity modulus, and the Poisson coefficient were assumed to be equal to those of the rachilla. The natural frequencies of the fruit-rachilla (eigenvalues), as well as the respective vibration methods (eigenvectors), were obtained through the formulation and calculation of the eigenvalue and eigenvector problems. For the modeling, systems with multiple degrees of freedom were considered, represented by a system of differential equations. The analysis of the dynamic behaviors was carried out using finite element discretization. The geometry discretization, modeling, and visualization of the results were conducted using the software Autodesk® Fusion 360®, student version.

**Table 1.** Mechanical, physical, and geometric properties used as input parameters during the computational modeling and simulation of the fruit-rachilla system for the accessions BD 27, BD 40, BGP 29, and BGP 35

|  | BD27 | BD 40 | BGP 29 | BGP 35 |
| --- | --- | --- | --- | --- |
| $d_r$ (mm) | **3.73** | **3.29** | **3.61** | **4.34** |
| $\rho_r$ (g cm<sup>-3</sup>) | **0.63** | **0.47** | **0.62** | **0.88** |
| a (mm) | **32.44** | **45.24** | **58.09** | **39.91** |
| b (mm) | **39.59** | **44.13** | **47.41** | **41.55** |
| c (mm) | **38.80** | **43.71** | **47.12** | **41.29** |
| $\rho_f$ (g cm<sup>-3</sup>) | **1.05** | **1.15** | **1.08** | **1.12** |
| E (MPa) | **238.90** | **260.70** | **293.90** | **194.00** |

| N | 0.37 | 0.37 | 0.37 | 0.37 |
| --- | --- | --- | --- | --- |

$d_r$ , rachilla diameter; $\rho_r$ , rachilla density; $a$ , largest characteristic dimension of the fruit; $b$ , intermediate characteristic dimension of the fruit; $c$ , lowest characteristic dimension of the fruit; $\rho_f$ , fruit density; $E$ , rachilla elasticity modulus, and $\nu$ , rachilla Poisson coefficient (source: Villar, 2016; Villar et al., 2017).

In the geometry discretization step, a convergence test was conducted to evaluate the type of mesh and the most adequate average element size. The convergence test yielded 10-node tetrahedral elements with an average side length of 0.001 m.

In the modeling step, three-dimensional models of the fruit-rachilla system were elaborated using the software Autodesk® Fusion 360®. The model was made up of a rachilla crimped at one of its extremities. The rachilla length were standardized as 12 cm. To simulate the system, the fruits were assumed to be attached at a position three-quarters of the rachilla length from the crimped side. The boundary conditions were also defined during the modeling step. Once the system modeling was defined, the Lanczos algorithm in the Autodesk® Fusion 360® software was used to determine the eigenvalues and eigenvectors.

Before the simulation, the model was validated using the natural frequencies acquired through the experiment process. A scanning test was conducted using ten samples from the fruit-rachilla system for each accession. To promote the excitation of the fruit-rachilla system, a Ling Dynamic Systems (LDS) instrument was used (Figure 1). The system was composed of a Dactron COMET_USB_ signal generator, an LDS PA 1000L amplifier, and an LDS V-555 electromagnetic vibrator. The scanning test involved increasing the frequency progressively by 1.5 octaves/min^-1^, from 10 to 40 Hz, with a peak-to-peak displacement amplitude of 0.5 mm. The system was excited with an acceleration rate 15-times the gravitational acceleration, and sampling at a rate of 500 Hz.

**Figure 1.**
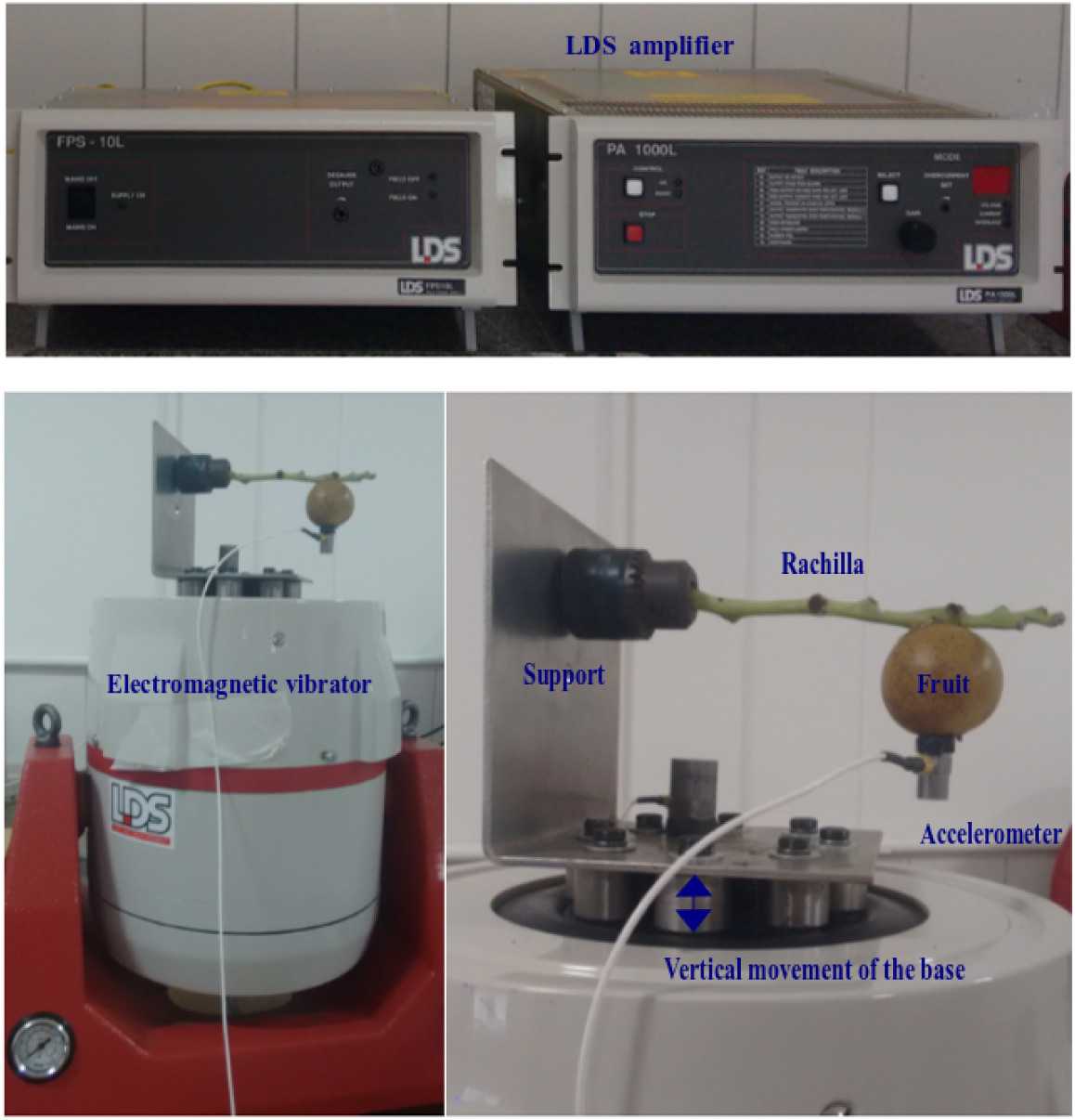
Data acquisition system for scanning test of fruit-rachilla system, composed of a Dactron COMETUSB signal generator, an LDS PA 1000L amplifier, an LDS V-555 electromagnetic vibrator and a piezoelectric accelerometer.

The fruit-rachilla samples were clamped at 2 cm lengthwise by using a fixed support in the vibrator, using the same length as in the geometric model. The sample fruits were attached to the rachilla at nearly the same place as for the geometric model. To acquire the data, a piezoelectric accelerometer LW174002 with a sensibility of 100.7 mVg^-1^, and an acquisition modulus cDAQ-9174 from National Instruments, were used. The accelerometer was fixed to the fruit for measuring the acceleration of the system in the vertical direction. The accelerometer mass was 5.6 g, which is 15 % less than that of the rachilla-fruit mass.

The acquired data were transformed into the frequency domain using a Fourier transform to find the resonance frequency that occurred at the peak displacement amplitude. This resonance occurred when the system excited frequency was equal to the fundamental frequency. To evaluate the models, the relative error between the measured and simulated fundamental frequencies was determined (Equation 1).

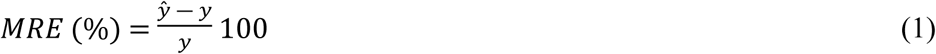

where,

MRE – mean relative error;

y_*i*_ – measured first natural frequency, Hz; and

ŷ_*i*_ – simulated first natural frequency, Hz.

A fundamental frequency was simulated using the average of the modulus of elasticity, fruit and rachilla density for all accessions. To identify which parameter had the greatest influence on fundamental frequencies, was carried out a sensitivity analysis. The rachilla’s modulus of elasticity and, the fruit and the rachilla density, were varied by ± 20 % on sensitivity analysis. Therefore, were obtained six fundamental frequencies and was compared with the average fundamental frequency.

## RESULTS AND DISCUSSION

The first natural frequencies were associated with resonance frequency. The resonance frequency was identified when occurred a peak amplitude displacement in frequency sampled range (Figure 2). The results presented in Table 2 list the mean relative errors and the descriptive statistical analysis of the first natural frequencies of each accession achieved experimentally through the scanning test and mathematically through the simulation. Some of the fruit-rachilla systems were eliminated because they presented frequencies above the pre-determined maximum limit of 40 Hz, which was attributed to errors occurring during the laboratory test.

**Table 2.**
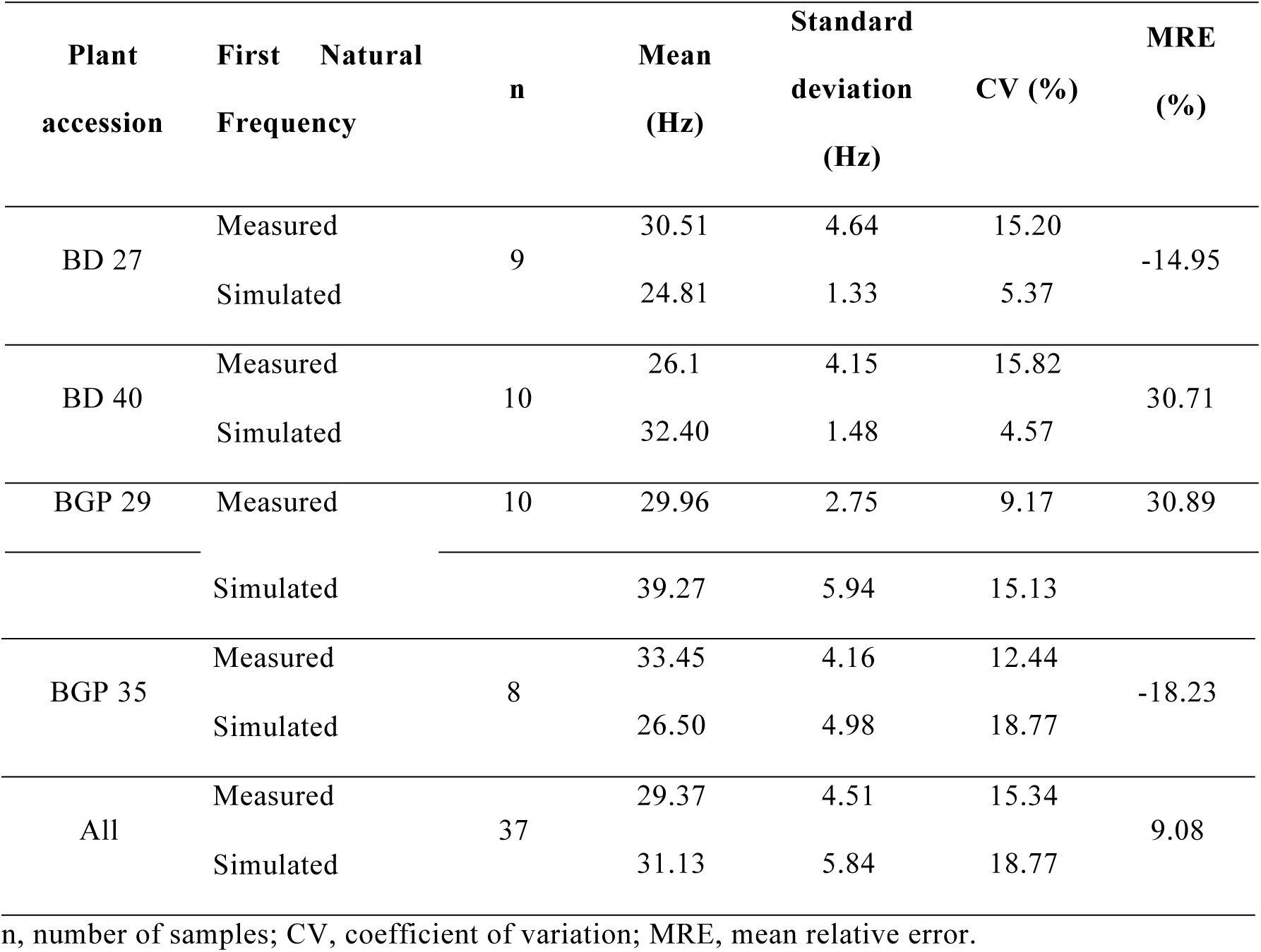
Descriptive statistics for the measured and simulated natural frequencies of the fruit-rachilla systems for the BD 27, BD 40, BGP 29, and e BGP 35 plants, as well as the mean relative errors between the measured and simulated values

**Figure 2.**
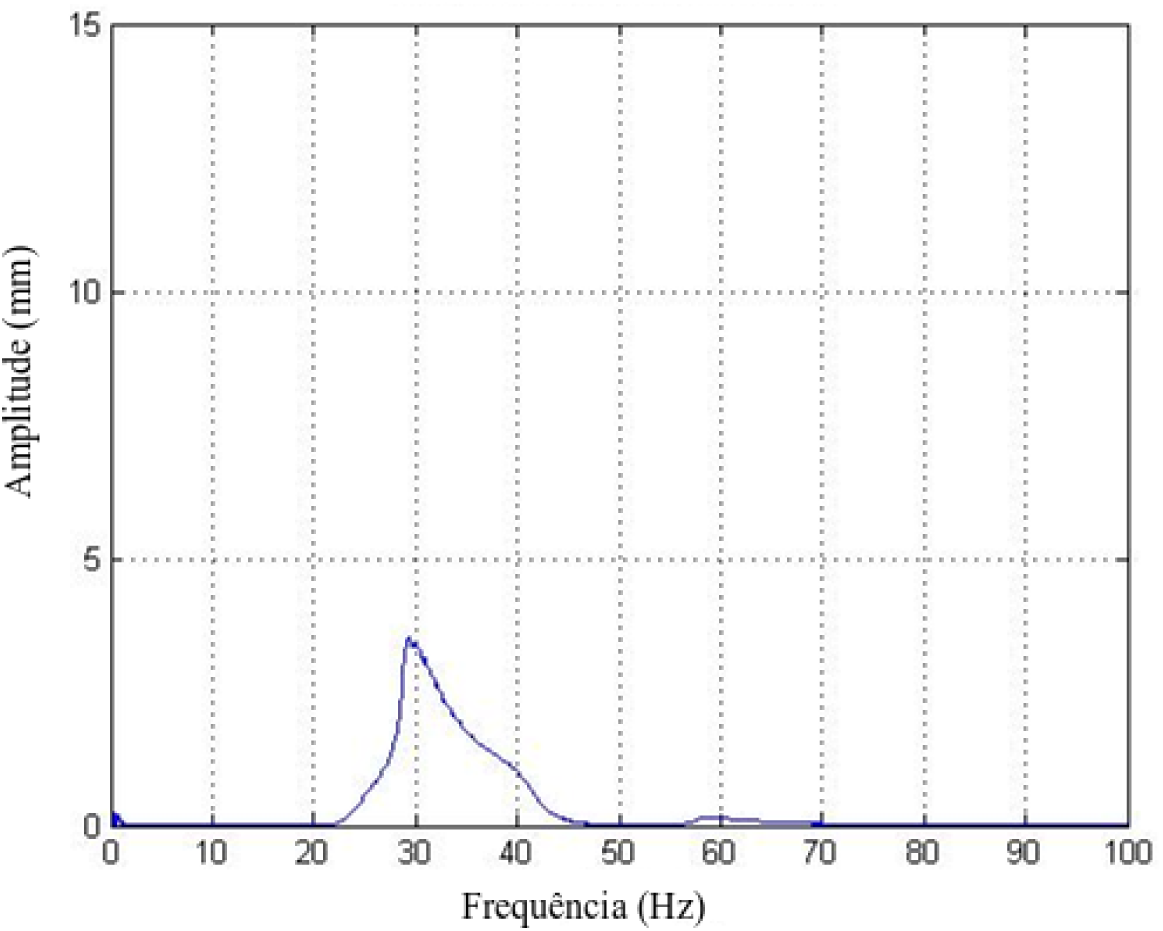
Frequency spectrum obtained using a Fourier transform to find the resonance frequency that occurred at the peak displacement amplitude.

**Figure 3.**
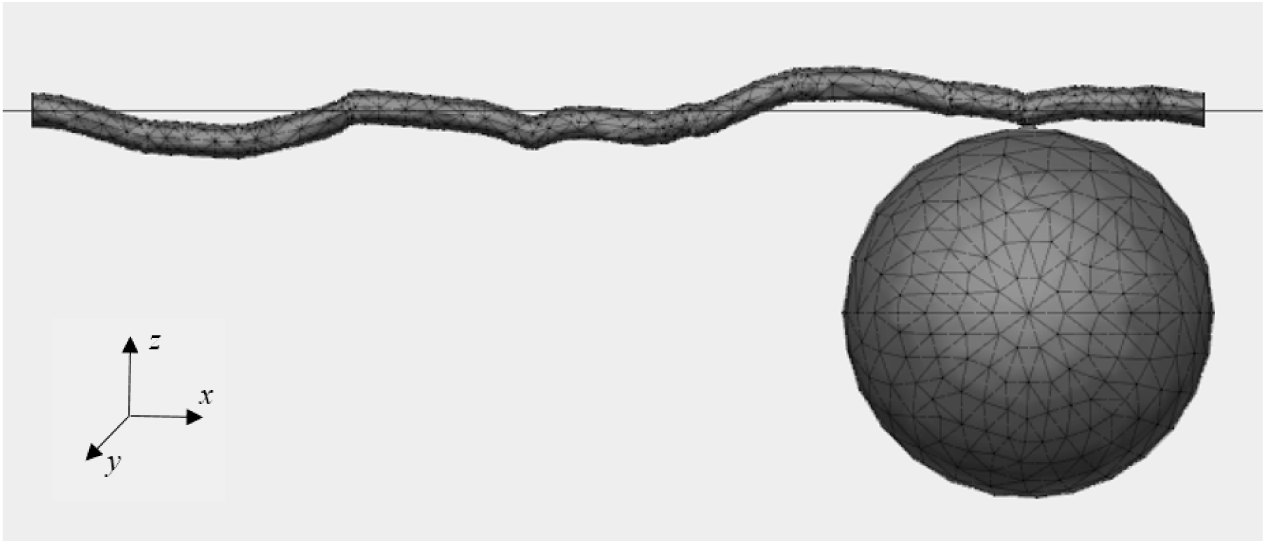
Fruit-rachilla model for Autodesk® Fusion 360® software finite-element simulation of modal analysis.

Because the accelerometer was positioned only for measurements in the z-direction, the modes of vibration obtained through the simulation for all plant accessions correspond to the vertical vibrations (Figure 1).

The natural frequencies obtained through the scanning test varied from 26.21 to 33.45 Hz on average with the coefficient of variation considered to be either low (between 0 % to 10 %) or medium (between 10 % to 20 %) for agricultural products (Gomes, 1987). The natural frequencies obtained through the mathematical simulation varied on average from 24.81 to 39.27 Hz with the coefficient of variation also considered as either low or medium for agricultural products. When analyzing the MRE of each plant accession, we observed that the proposed mathematical model presented a significant variation in estimating the natural frequencies with an MRE ranging from -14.95 to 30.89%.

The best result obtained during the simulation was for accession BD 27 with an MRE equal to -14.95%. When the MRE is closer to zero, the model’s capacity to estimate the natural frequencies is better accuracy. The worst result was for accession BGP 29 with an MRE equal to 30.89%. The model underestimated the fundamental frequency in two plants accessions (negative MRE) and overestimated (positive MRE) in two other plants accessions. Thus, the model did not show trend by underestimating or overestimating the experimental data. This behavior, most likely, due to mechanical and geometrical properties variation between plants of different accessions.

Santos (2008) conducted a modal analysis for the fruit-peduncle system of a coffee plant using the finite element method and compared the results with an analytical solution and achieved a maximum MRE of 1.88%. In the present study, an analytical solution was not obtained. However, was carried out a comparison of the values which were obtained experimentally, with values obtained by simulation. The determined MRE can be explained through the approaches and considerations applied to enable the simulation.

The elasticity modulus of the fruits, and the ratio Poisson, used as input parameters during the simulation were the same as those applied for the rachilla, and the rachilla was considered with a unique diameter. These considerations may have influenced the model results because the actual rachilla diameter is not constant, and the fruits likely present mechanical properties differing from those of the rachilla.

The first natural frequencies were influenced by the elasticity modulus and fruit density (Table 3). When density of fruit and elasticity modulus were decreased by 20%, the fundamental frequency was increasing by 11.79% and decreasing by 4.69% on average, respectively. When density fruit and elasticity modulus were increased by 20%, the fundamental frequency was decreasing by 8.69% and increasing by 4.07% on average, respectively.

**Table 3.**
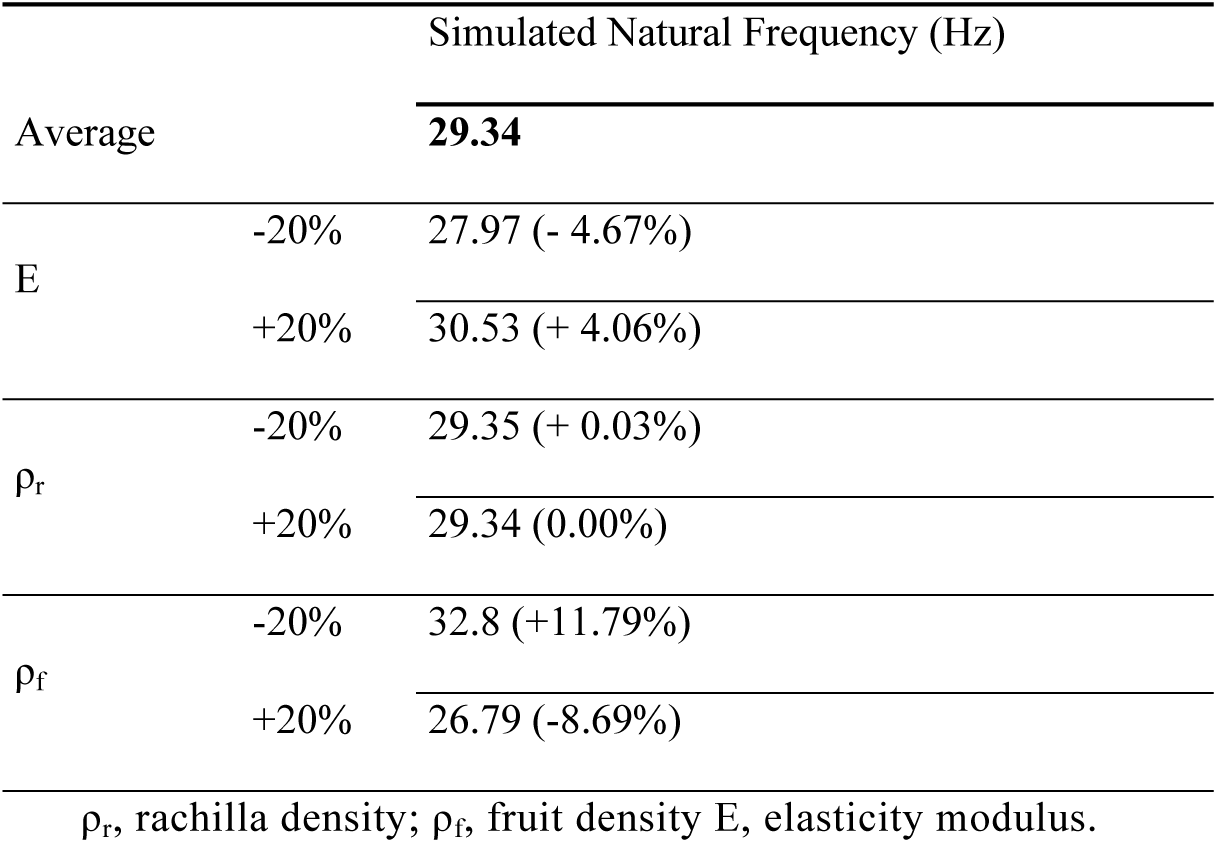
Average simulated natural frequencies during the sensibility analysis when the density, elasticity modulus, and Poisson coefficient were varied by ± 20%, and the differences between the average simulated natural frequency of the model (as a percentage)

The density of rachilla property showed to have the lowest effect on the first natural frequency. The first natural frequency decreases 0.03% when the density of rachilla was increased by 20%. No variation in natural frequency was observed when the density of rachilla was decreased by 20%. The natural frequency is directly proportional to the hardness, as determined based on the geometry and elasticity modulus, and inversely proportional to the mass, which is related to the density (Rao, 2008). Therefore, when an increase in the elasticity modulus or a decrease in the density occurs, the natural frequency is higher, as was seen in the analyzed case, showing a positive difference.

## CONCLUSION

- The model used to estimate the modal properties of the macaw palm fruit-rachilla system (natural frequencies and vibration modes) was development and showed mean relative error of 9.08%.
- T The first natural frequencies obtained were 29.37 and 31.13 Hz, for experimental and simulated data, respectively.
- T The parameters showing a greater influence on the estimation of the natural frequencies were the elasticity modulus and the density of fruits.
- T The natural frequencies determined are associated with the vertical vibrations.

## ACKNOWLEDGEMENT

The authors of this work wish to thank FAPEMIG (Fundação de Amparo à Pesquisa do Estado de Minas Gerais), CAPES (Coordenação de Aperfeiçoamento de Pessoal de Nível Superior) and CNPq (Conselho Nacional de Desenvolvimento Científico e Tecnológico) for their financial support.

